# Rapid population flux in bacterial spot xanthomonads during a transition in dominance between two genotypes in consecutive tomato production seasons and identification of a new species *Xanthomonas oklahomensis* sp. nov

**DOI:** 10.1101/2025.04.13.648550

**Authors:** Brett Johnson, Aastha Subedi, John Damicone, Erica M. Goss, Jeffrey B. Jones, Mustafa O. Jibrin

## Abstract

In the bacterial spot of tomato disease complex, *Xanthomonas euvesicatoria* pv. *perforans* (*Xep*) is known to outcompete *X. euvesicatoria* pv. *euvesicatoria* (*Xee*). The result, over time in tomato production systems where both pathogens are present, is the anticipated displacement of *Xee* by *Xep*. In this study, we characterized the population of recovered strains from grower fields in response to a bacterial spot of tomato outbreak in east and central Oklahoma in 2018 and 2019. Tomato and pepper strains sampled in earlier years were included to provide additional context. Phenotypic and genome-based analyses showed marked differences in race and species composition in recovered strains. All pre-2018 (2001-2014) tomato bacterial spot strains were *Xep* race T3, except one T4 strain. Tomato bacterial spot strains in 2018 consisted of *Xee* race T1 and *Xep* T3, while strains recovered in 2019 from the same locations were exclusively *Xep* T4 strains. The 2019 *Xep* T4 strains form the same cluster with the 2014 *Xep* T4 strain on the phylogenetic tree, suggesting the same population. Only the *Xep* race T4 strains showed an expanded host range to pepper. The recovered strains were also variable in copper sensitivity and effector content. We additionally recovered non-bacterial spot xanthomonads, one of which belongs to the new species, *Xanthomonas oklahomensis* sp. nov. These results contributes to novel insights in understanding genomic heterogeneity and seasonal population flux between competing bacterial genotypes during disease outbreaks and can be considered when developing disease management strategies.

## Introduction

Plant pathogen populations in agronomic systems are continuously under a state of flux, with the population changing at different rates depending on the impact of factors such as the disease state of the planting material, type of the pathogens already present in a given location, migration of new strains into the location, interactions within and between the microbiome, local production and management practices as well as climatic conditions (McDonald and Stukenbrock, 2016; Klein-Gordon *et al*. 2021; Zhan *et al*. 2022; Laine, 2023; Singh *et al*. 2023; Lahlali *et al*. 2024; Timilsina *et al*. 2025). In bacterial species, clonal reproduction is expected to maintain clonal lineages. However, homologous recombination and horizontal gene transfer, alongside migration, increases the genetic diversity (Jones, 2021; Montgomery *et al*. 2023). These processes have short- and long-term consequences on population structure and species composition (Treindl *et al*. 2023; Sharma *et al*. 2024;Gilbert and Parker, 2010; Rahnama *et al*. 2023).

Population changes over a long period of time are expected norms for plant pathogens in agronomic systems (Yoshida *et al*. 2013; Goss *et al*. 2014; Coomber *et al*. 2024; Jibrin *et al*. 2024; Timilsina *et al*. 2025). However, plant associated bacterial species experience rapid and short-term population changes that have consequences on existing population dynamics. Rapid changes in pathogen populations over a short term can add to an already difficult terrain of bacterial disease management, especially where these population changes have consequences for host-pathogen interactions and chemical-based management (Bonneaud and Longdon, 2020; Ristaino *et al*. 2021; Fielder *et al*. 2024).

Within the bacterial spot of tomato and pepper disease pathogen complex, various levels of population change that impact pathogen epidemiology and disease management have been uncovered. Bacterial spot disease is caused by four genetically distinct groups with highly variable genetic structure belonging to the genus *Xanthomonas*. These include *X. euvesicatoria* pv. *euvesicatoria* (*Xee*, Syn. *X. euvesicatoria*)*, X. euvesicatoria* pv. *perforans* (*Xep*, Syn. *X. perforans*)*, X. hortorum* pv. *gardneri* and *X. vesicatoria* (Jones *et al*. 2004; Osdaghi *et al*. 2021; Jibrin *et al*. 2022). Coexistence between combinations of the four genotypes is not uncommon, and studies have additionally identified sub-groups and sub-populations that are indicative of unique evolutionary events within each species (Kebede *et al*. 2014; Timilsina *et al*. 2015; Jibrin *et al*. 2018; Newberry *et al*. 2019; Roach *et al*. 2019; Timilsina *et al*. 2019; Chen *et al*. 2024; Jibrin *et al*. 2024, Subedi *et al*. 2024a; Timilsina *et al*. 2025). Alongside disease epidemics favored by warm and humid conditions, rapid change in the population of the bacterial spot pathogen makes the disease especially difficult to control (Osdaghi *et al*. 2021; Jibrin *et al*. 2022).

In the continuous monitoring of bacterial spot disease in tomato and pepper production systems in Florida, understanding changes in bacterial spot pathogen populations have been critical to understanding pathogen epidemiology and disease management. It is now known that the previously prevalent *Xee* strains have been displaced by *Xep* in tomato fields in Florida within a 15-year period (Tudor-Nelson *et al*. 2003; Hert *et al*. 2005; Hovarth *et al*. 2006; Timilsina *et al*. 2019; Klein-Gordon *et al*. 2021). Strains of *Xep* produce bacteriocins against strains of *Xee* and were likely responsible for the displacement of *Xee* in Florida tomato fields (Tudor-Nelson *et al*. 2003; Hert *et al*. 2005). Additionally, widely prevalent copper resistant *Xep* strains as well as non-durability of resistance genes have made management of bacterial spot in tomato production systems difficult in Florida (Stall *et al*. 2009; Timilsina *et al*. 2019; Bibi *et al*. 2024; Kaur *et al*. 2024). In the Taiwan tomato production system too, *Xee* was the dominant pathogen between 1989-2003 (Chen *et al*. 2024). Post 2003-2009, *Xep* replaced *Xee* as the bacterial spot pathogen in Taiwan in a period of about eight years (Chen *et al*. 2024). The prevalence of *Xep* in tomato production systems has recently been recorded in multiple production systems (Araujo *et al*. 2017; Egel *et al*. 2018; Jibrin *et al*. 2018; Adhikari *et al*. 2019).

The objective of this study was to characterize recovered bacterial spot strains following an outbreak of bacterial spot disease in the tomato production system in Oklahoma during the 2018 and 2019 growing seasons to understand species composition and diversity. The majority of strains used in this study were collected from a 2018 and 2019 epidemic in central and eastern Oklahoma. A few strains from previous years were added to provide context. Our results are important in understanding potential seasonal population changes between competing bacterial genotypes and implications for disease management.

## Materials and methods

### Survey and bacterial isolation

Field surveys were carried out during the 2018 and 2019 tomato production season in Oklahoma in 29 field sites across 13 counties in Central and Eastern Oklahoma. Sites were selected based on reports by growers or extension personnel of the presence of an unidentified foliar disease in field-grown tomatoes. Tomato fields in Rogers, Cherokee, and Bryan counties had Septoria leaf spot and early blight, but no bacterial spot diseases, and were not included in this study. Figure 1 shows counties where field sites had confirmed tomato bacterial spot disease in 2018 and 2019. Samples were collected from symptomatic plants, transported in plastic bags, and kept at 4°C until isolation. Bacteria were isolated from lesions consistent with symptoms of bacterial spot. Lesions were excised into 2 mm^2^ sections, and surface sterilized in 0.05 % sodium hypochlorite and 10 % ethyl alcohol for 30 s. Sections were rinsed for 60 s in sterile distilled water, cut into smaller pieces in a drop of sterile distilled water, incubated for 30 min, and streaked to nutrient agar (NA). Cultures were incubated at 28°C for 48 h and yellow-mucoid colonies typical of *Xanthomonas* species were re-streaked to obtain single colonies. Pure single colonies were re-streaked to obtain a lawn of bacterial growth on plates which was subsequently stored in 15% glycerol at -70 °C. A total of 24 isolates that represented 24 field samples were collected in 2018. A total of 42 isolates that represented 24 field samples were collected in 2019.

**Figure 1.**
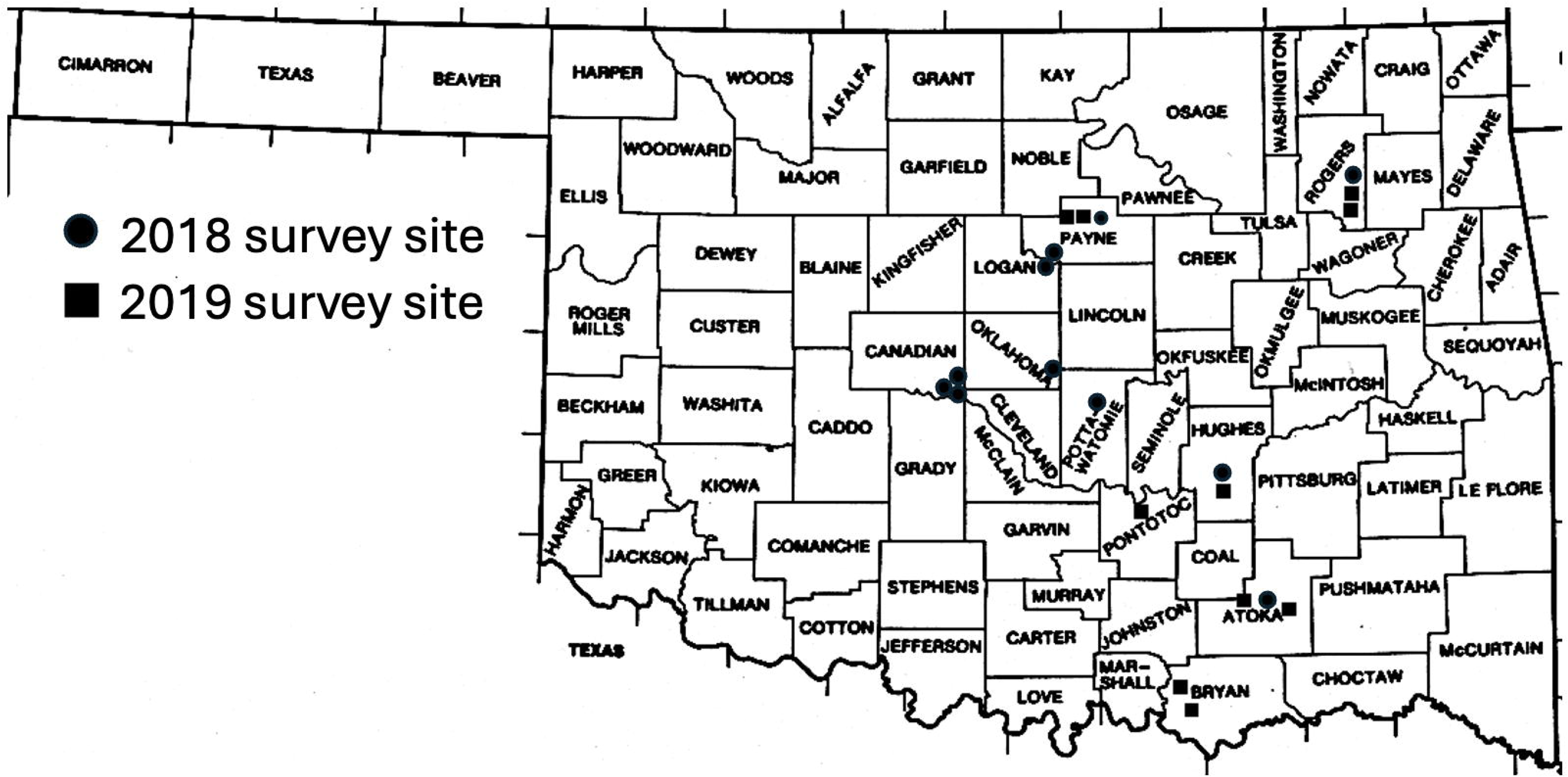
Map of Counties in Oklahoma where field sites had confirmed bacterial spot in 2018 (n=11 sites) and 2019 (n=10 sites).

### Preliminary screening of recovered isolates

The pathogenicity of all strains was assessed by infiltrating bacterial suspensions (adjusted to 10^8^ CFU/ml, OD600 = 0.3) from fresh overnight cultures into pepper cultivar ‘Early California Wonder’ (ECW) and tomato cultivar ‘Bonny Best’. Strains that did not elicit a hypersensitive response (HR) within 24-48 hr but caused necrosis within 72 to 96 hr were recorded as pathogenic. A set of 59 isolates, representing isolates collected from 1998 – 2019 and 4 reference isolates previously identified as pathogenic on tomato were identified using RST65/RST69 PCR primers that amplify a 420 bp fragment of the *hrcN* gene conserved among bacterial spot xanthomonads (Obradovic *et al*., 2004). Isolates were grown on nutrient agar for 48 h at 28 °C. DNA was extracted using DNeasy Ultra Clean Microbial Kit (QIAGEN, Germantown, MD) and suspended in TE buffer. All DNA samples were diluted to a concentration of 20 ng/uL for use in PCR reactions. The PCR master mix consisted of 2 uL of DNA, 5.5 uL of sterile nuclease free water, 2.5 uL of each of two primers (RST65, RST69), and 12.5 uL of Taq polymerase. PCR amplification was conducted in the thermocycler under the following conditions: 1 cycle of 95 °C for 5 m, 29 cycles of 95 °C for 30 s, 63 °C for 1 m, 72 °C for 45 s, followed by 1 cycle 72 °C for 5 m. The PCR products were purified using illustra GFX PCR DNA and Gel Band Purification Kit (GE Life Sciences, Marlborough, MA). Amplicons were Sanger sequenced in the Core Facility of the Department of Biochemistry and Molecular Biology at Oklahoma State University. Forward and reverse sequences of sequenced DNA were assembled using BioEdit (https://thalljiscience.github.io/) and compared to sequences on the NCBI database using BLAST. Assembled sequences were deposited on NCBI database.

### Pathogenicity and race determination

Based on the results of preliminary screening, a total of 27 strains including strains isolated during the 2018 and 2019 surveys, and a few strains isolated between 1998 and 2014 were selected for further studies. Identification of races of these strains was carried out on pepper and tomato as described previously (Bouzar *et al*. 1994; Stall *et al*. 2009). Briefly, overnight cultures of tomato strains were infiltrated into leaves of tomato genotypes Hawaii 7998, FL 216 and LA716 harboring *rxv, Xv3* and *XopJ4/AvrXv4* resistance genes, respectively. *X. euvesicatoria* E3-1 and *X. perforans* 91-118 were used as control strains (Klein-Gordon *et al*. 2021). Similarly, bacterial suspensions of strains isolated from pepper were infiltrated into three near-isogenic lines of ECW carrying single resistance genes *Bs1* (ECW10R), *Bs2* (ECW20R), and *Bs3* (ECW30R). Strains Xe_(82-8 UNS::pXvCu), Xe_88-5, and Xe_95-2 were used as positive controls for incompatible reactions with *Bs1, Bs2,* and *Bs3,* respectively (Subedi *et al*. 2023). Infiltrated leaf areas were monitored for HR or susceptible reactions at 12, 24-, 36-, 48-, and 72-hr post-inoculation on both tomato and pepper genotypes.

### Copper and streptomycin sensitivity and amylase activity

The 27 strains were evaluated for copper and streptomycin sensitivity, and amylase activity following the methods described previously (Subedi *et al*. 2023). For the copper sensitivity assay, strains were cultured overnight on NA amended with 20 µg/ml of copper sulfate pentahydrate at 28°C to induce copper resistance genes. Bacterial cells from these cultures were picked with a toothpick and placed on NA amended with 200 µg/ml copper sulfate pentahydrate. For determining streptomycin sensitivity and amylase activity, 24-hr fresh cultures grown on NA were transferred onto NA amended with streptomycin (200 µg/ml) or soluble starch (1.5%), respectively (Jones *et al*. 1995; Stall *et al*. 1994). Plates were incubated at 28°C and evaluated for 48 hr. Copper and streptomycin tolerance were considered positive by observing visible bacterial growth on amended media, while a turbid halo around each colony indicated positive amylase activity. *X. euvesicatoria* E3-1 and *X. euvesicatoria* 71-21 were used as positive and negative control strains for copper and streptomycin sensitivity, respectively, and *X. euvesicatoria* E3-1 and *X. perforans* 91-118 were used as positive and negative controls for amylase activity. Each strain was tested in two replicates.

### DNA extraction, genome sequencing and assembly

Pure cultures of the 27 selected strains were streaked on NA plates and incubated at 28°C. After 24 hr, the bacterial colonies were transferred to test tubes containing nutrient broth and incubated overnight at 28°C while shaking at 250 rpm. Genomic DNA was extracted from these overnight cultures using the gram-negative bacterial DNA extraction protocol provided by the Wizard Genomic DNA Purification Kit (Promega, Madison, WI). The concentration of the extracted DNA was quantified using a Nanodrop spectrophotometer (Thermo Fisher Scientific Inc., Wilmington, DE). Whole-genome sequencing of DNA samples was carried on the Illumina NextSeq 2000 platform by the Seq Center (Pittsburgh, PA, USA), resulting in 151 bp paired-end reads. The genome assembly of the raw sequences was carried out as previously described (Timilsina *et al*. 2019). Briefly, Trim Galore was used to remove adapters, clean the sequences, and pair the raw reads. The paired reads were then assembled using SPAdes (v. 3.10.1), and contigs shorter than 500 bp with K-mer coverage less than 2.0 were filtered out (Nurk *et al*., 2013). The validated reads were aligned to the filtered contigs using Bowtie 2 (v. 2.3.3) (Langmead and Salzberg, 2012), generating SAM files that were converted to BAM files using SAMtools (Li *et al*., 2009). Pilon was utilized to polish and refine the draft genome assemblies (Walker *et al*., 2014). Genome statistics were calculated using a custom Python script available in https://github.com/sujan8765/nepgorkhey_python/blob/master/genome_stats.py. Prokka (v. 1.10) was used to annotate the assembled genomes with default parameters (Seemann, 2014).

### Taxonomic analyses

The assembled genomes were submitted to the Type Strain Genome Server (TYGS) (https://tygs.dsmz.de/) for strain identification and taxonomic analyses using described methods (Meier-Kolthoff and Goker, 2019). Using the closest type strains based on results from TYGS, whole genome pairwise average nucleotide identity (ANI) was calculated among the assemblies and compared to the reference genomes of type strains *X. euvesicatoria* ATCC 11633, *X. perforans* Xp-DSM-18975, *X. populi* CFBP 1817, *Xanthomonas cynarae* CFBP 4188, *X. gardneri* ATCC 19865, *X. hortorum* CFBP 4925, *X. hydrangeae* GBBC 2123T, *X. arboricola* CFBP 2528, and *X. alfalfae subsp. citrumelonis* CFBP 3371 using pyani (v. 0.2.10) (Pritchard *et al*., 2016). Digital DNA-DNA hybridization (dDDH) values and confidence intervals were calculated using the recommended settings in GGDC 4.0 on TYGS (Meier-Kolthoff *et al*. 2022). Proksee (https://proksee.ca; Grant *et al*. 2023) was used to visualize the circular genome of the potentially new species.

### Core genome phylogenetic analysis

Core genes of sequenced strains in this study, alongside previously sequenced strains obtained from NCBI database, were identified using Roary (v. 3.12.0) with a minimum BLASTp identity of 95% (Page *et al*., 2015). The nucleotide alignment of core genes was conducted using MAFFT (Katoh and Standley, 2013). Phylogenetic analysis was carried out on the core genome alignment output from Roary using RaxML (Stamatakis, 2014) under the GTR GAMMA I substitution model. The best-scoring tree from RasxML was then processed with ClonalFrameML to produce a recombination-corrected tree (Didelot and Wilson, 2015). The resulting tree was visualized using iTOL (v. 5.6.3) (Letunic and Bork, 2007).

### Type III Secretion Effector Analysis (T3SEs) and presence of copper resistance and amylase genes

A database of 64 known Type III secretion effector (T3SE) proteins from various xanthomonads (Parajuli *et al*. 2024), was used to screen for the presence of T3SEs using tBLASTn with an E value threshold greater than 10^-5^. Effector presence was defined by using the criterion of over 75% amino acid identity covering more than 75% of the query length. This analysis focused solely on the presence or absence of effectors based on the specified thresholds and did not account for mutations or insertion elements that could affect protein functionality. For TAL effectors, only TALE hits were reported due to the inability of short reads to accurately assemble their repetitive regions. A similar approach was used to screen genomes for the presence of copper resistance genes, *copLAB* and the amylase gene (Subedi *et al*. 2024b).

## Results Survey results

Bacterial spot disease was identified in growers’ tomato fields in seven counties in 2018 and six counties in 2019 (Table S1.1, Supplementary File 1,). The impact of the disease, as measured by disease incidence on the leaves and whole plant as well as defoliation, was a nearly 2-fold increase in 2019 compared to 2018.

Preliminary identification based on the partial sequences of the *hrcN* gene is presented in Tables S1.2 and S1.3 (Supplementary File 1). Results of the partial sequences of the *hrcN* gene identified pre-2018 tomato strains as belonging mostly to *Xee* and *Xep*, with one potentially *X. euroxanthea* strain and one *X. vesicatoria* strain; 2018 strains similarly contained a mixture of *Xee* and *Xep,* including one potentially *X. arboricola* pv. *pruni;* while the majority of 2019 strains were *Xep*, with one potentially *X. arboricola* pv. *juglandis* strain.

### Pathogenicity and race distribution

All tomato *Xee* and *Xep* strains were pathogenic on tomato cultivar Bonny Best while the pepper strains were pathogenic on pepper only (Table S2.1, Supplementary File 2). Within the *Xep* strains, 16 tomato strains showed susceptible reactions on pepper cultivar ECW (Table S2.1, Supplementary File 2). All strains isolated from tomato, except strains XCV_T1, XCV_T19_1 and XCV_T19_11 showed susceptible reactions on Bonny Best while all 4 strains isolated from pepper elicited an HR on Bonny Best but susceptible reactions on the pepper line ECW. For the strains that were pathogenic on tomato, tomato races T1 (5 strains), T3 (8 strains), and T4 (7 strains) were identified (Table S2.1, Supplementary File 2). Strains isolated from pepper were P1 races (Table S2.1, Supplementary File 2). The reactions of strains XCV_T1, XCV_T19_1 and XCV_T19_11 on tomato and pepper differential lines did not fit into known described race classifications.

### Copper and Streptomycin sensitivity, Amylase activity

The results of copper and streptomycin sensitivity, and amylase activity are presented in Table S2.1 (Supplementary File 2). All T1 and P1 strains, along with three T3 and four T4 strains were sensitive to copper. However, five T3 strains and three T4 strains were tolerant to copper. Some *Xep* strains were tolerant to copper, whereas all *Xee* strains were sensitive to copper. All strains were sensitive to streptomycin. All *Xep* strains were strongly amylolytic. Approximately 50% of the *Xee* strains were copper tolerant, 5 *Xee* tomato strains were amylase positive while the 4 *Xee* pepper strains were non-amylolytic. All *Xep* T3 and T4 strains were strongly amylolytic. Strains XCV_T1, XCV_T19_1 and XCV_T19_11 were also amylase positive.

### Genome sequencing information, average nucleotide identity and *in silico* (is-) DDH analyses

Genome sequencing information of all sequenced strains along with the NCBI accessions are presented in Table S2.2 (Supplementary File 2). Preliminary determination of closest type strains using a complementary 16S rDNA and genome BLAST distance phylogeny approach methods on the Type Strain Genome Server (TYGS) confirmed that majority of the strains belong to *Xee* and *Xep* (Fig. S1 and Fig. S2, Supplementary File 3). However, the three strains that elicited HRs in tomato and pepper grouped differently. One strain, XCV_T19_11, clustered with the type strain, *Xanthomonas alfalfae* subsp. *citrumelonis* CFBP 3371, the causal organism of citrus bacterial spot.; the second strain, XCV_T19_1, clustered with the type strain of *Xanthomonas arboricola* CFBP 2528, which causes walnut blight; and the third strain, XCV_T1, was unique, suggesting a potentially new species. Average nucleotide identity comparisons provided similar results to TYGS analyses. Based on the average nucleotide identity (ANI) with type strains from different *Xanthomonas* species, the genomes of nine strains shared >99.7 % identity with the type strain *X. euvesicatoria* ATCC 11633 while the genomes of 16 strains shared >99.7 % identity with type strain of *X. perforans* DSM_18975 (Table S2.3, Supplementary File 2). Strains XCV_T19_1 and XCV_T19_11 have more than 98.7% and 99.7% ANI with type strains *Xanthomonas arboricola* CFBP 2528 and *Xanthomonas alfalfae* subsp. *citrumelonis* CFBP 3371, respectively. The TYGS predicted XCV_T1 strain as a potentially new species, which was closest to *X. hydrangeae* LMG 31884 based on GBDP and by ANI with 91% similarity (Table S2.3, Supplementary File 2; Fig S3, Supplementary File 3). Comparisons based on the digital *in silico* DNA-DNA hybridization (is-DDH) further confirmed that this strain belongs to a new species (Table 3; S2.4). Based on is-DDH the closest strains to the XCV_T1 based on is-DDH are *X. cynarae* CFBP4188 and *X. hydrangea* with is-DDHs of 67.6% and 66.2%, respectively. We have named this new species *Xanthomonas oklahomensis* sp. nov., with the newly assigned NCBI taxonomy ID, 3391446.

#### Phylogenetic analysis based on core genome

The core genome phylogenetic tree of all sequenced strains along with a subset of previously sequenced strains showed a similar result to ANI (Fig. 2a). XCV_T19_1 was closest to *Xanthomonas arboricola* CFBP 2528 and XCV_T19_11 to *Xanthomonas alfalfae* subsp. *citrumelonis* CFBP 3371 (Fig. 2a). The *Xee* strains isolated from tomato in Oklahoma formed a single clade different from the *Xee* strains isolated from pepper (Fig. 2b). The *Xee* tomato strains grouped in previously identified clade of strains with an amylolytic phenotype, which includes strains from Florida, Ohio, Mexico, Taiwan (Fig. 2b; Subedi *et al*. 2024b). The *Xee* pepper strains cluster closely with strain Tu10 from Turkey (Fig. 2b, Subedi *et al*. 2024b). The amylolytic tomato *Xee* strains from Oklahoma clustered with the amylolytic *Xee* strains from Florida (19_200E2, 20_26_K2, 20_21_R2), Ohio (OH_1605), Mexico (Mex_1080), Taiwan (XVP-312 and XVP-318), with an intact alpha-amylase gene (Fig. 2b; Subedi *et al*. 2024b). However, Oklahoma pepper *Xee* strains harbor the frameshift deletion in the alpha amylase gene reported in non-amylolytic *Xee* strains and grouped with non-amylolytic strains (Fig. 2b; Barak *et al*. 2016; Subedi *et al*. 2024b). All 2019 *Xep* strains isolated from tomato, plus a 2014 strain (XCV14_T3), grouped with strains that were recently identified to show expanded host range to pepper (Fig. 2c; Subedi *et al*. 2024a). All other pre-2019 *Xep* strains clustered separately from the 2019 strains (Fig. 2c).

**Figure 2.**
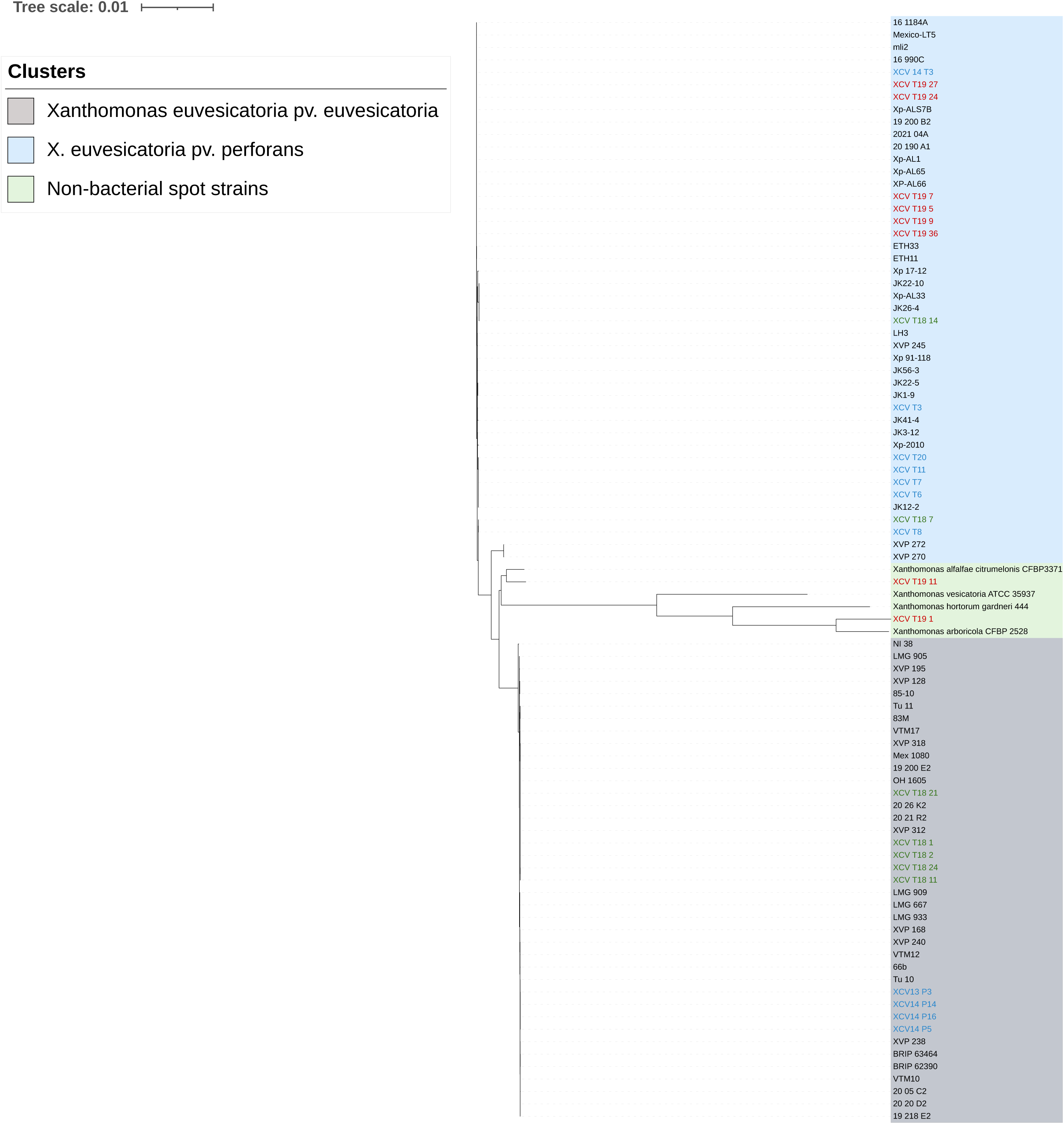
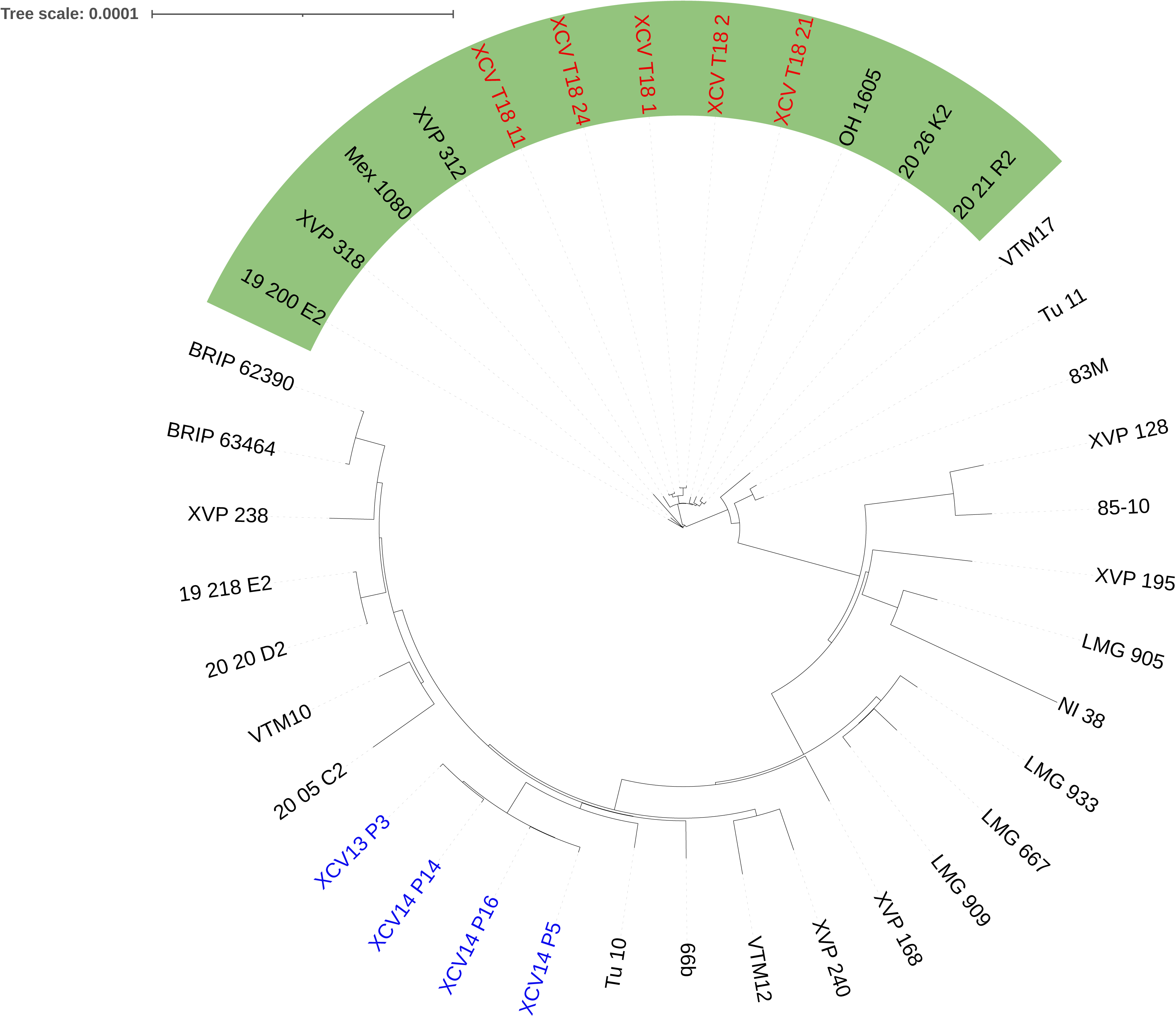
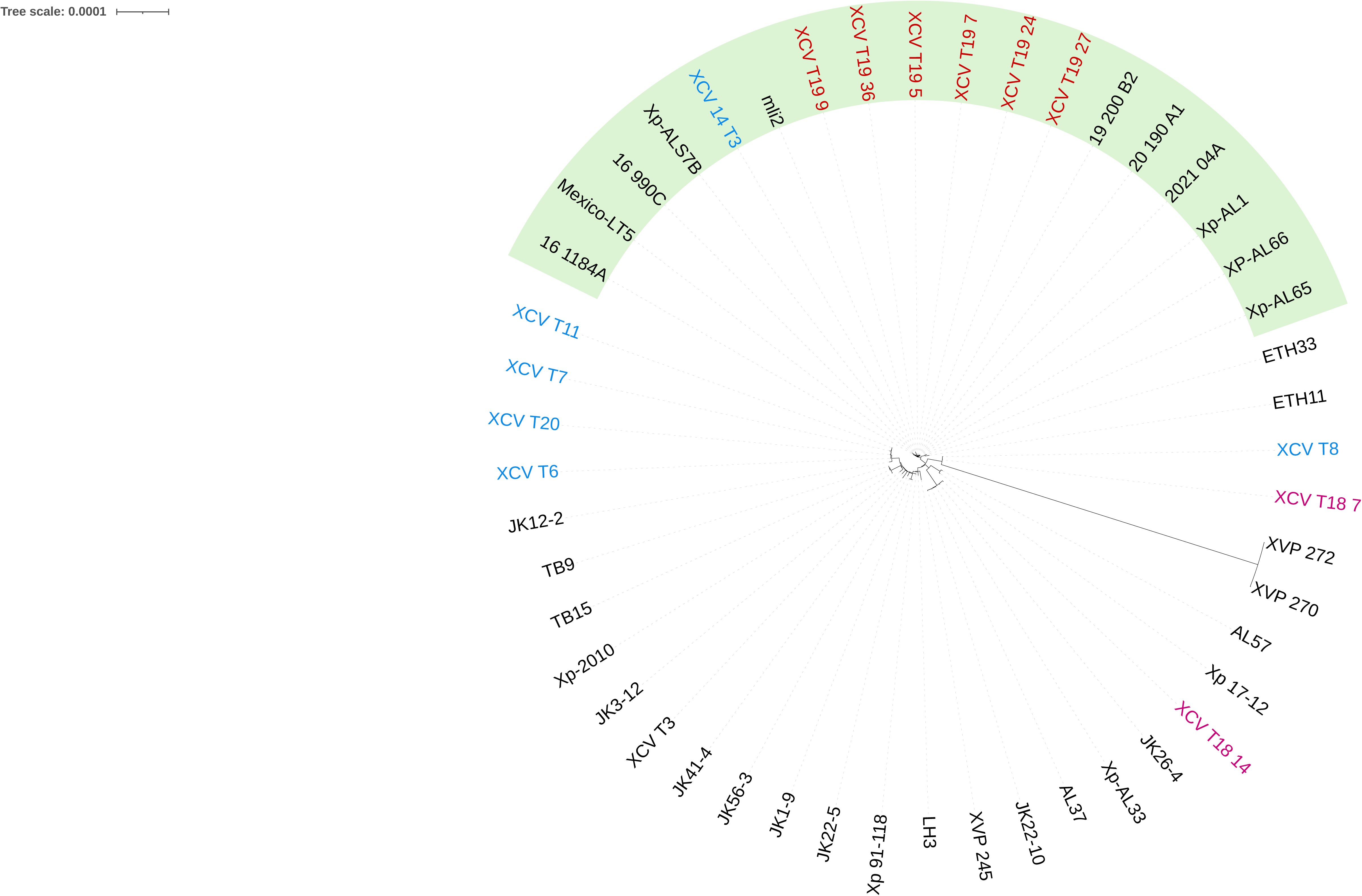
(a) Maximum likelihood phylogeny based on the shared genes between the genomes of Oklahoma strains compared with genomes of previously sequenced strains from NCBI database. (b) Maximum likelihood phylogeny based on the shared genes between the genomes of Oklahoma *Xanthomonas euvesicatoria* pv. *euvesicatoria* (*Xee*) strains and other *Xee* strains from NCBI database. (c) Maximum likelihood phylogeny based on the shared genes between the genomes of Oklahoma *Xanthomonas euvesicatoria* pv. *perforans* (*Xep*) strains and other *Xep* strains from NCBI database. For all figures, strains in blue are pre-2018 Oklahoma strains, strains in green are 2018 Oklahoma strains while strains isolated in 2019 are in red. Amylolytic *Xee* strains are shown in green background in Figure 2b. *Xep* strains with expanded host range to pepper are shown in green background in Figure 2c. Strain XCV_T1 was not included in the tree.

### Type 3 Effector Profiles

Out of 64 effectors investigated, 48 effectors had presence/absence variability based on the cutoff of >75% identity and coverage in 27 sequenced strains (Fig. 3). All *Xee* and *Xep* strains possessed the previously identified core effector genes found in bacterial spot xanthomonads (Potnis *et al*. 2011). These core effectors include AvrBs2, XopD, XopF1, XopK, XopL, XopN, XopQ, XopR, XopX, XopZ1, XopAD. Additionally, *XopAU*, which was originally characterized in *Xee*, was present in all the sequenced *Xee* and *Xep* strains. The effector AvrBsT, was only found in pre-2018 strains. All tomato and pepper *Xee* strains had hits for a TAL effector. However, none of the *Xep* strains contained TAL effectors. In general, thirty effectors were considered common effectors, defined as present in more than 50% of the strains.

**Figure 3.**
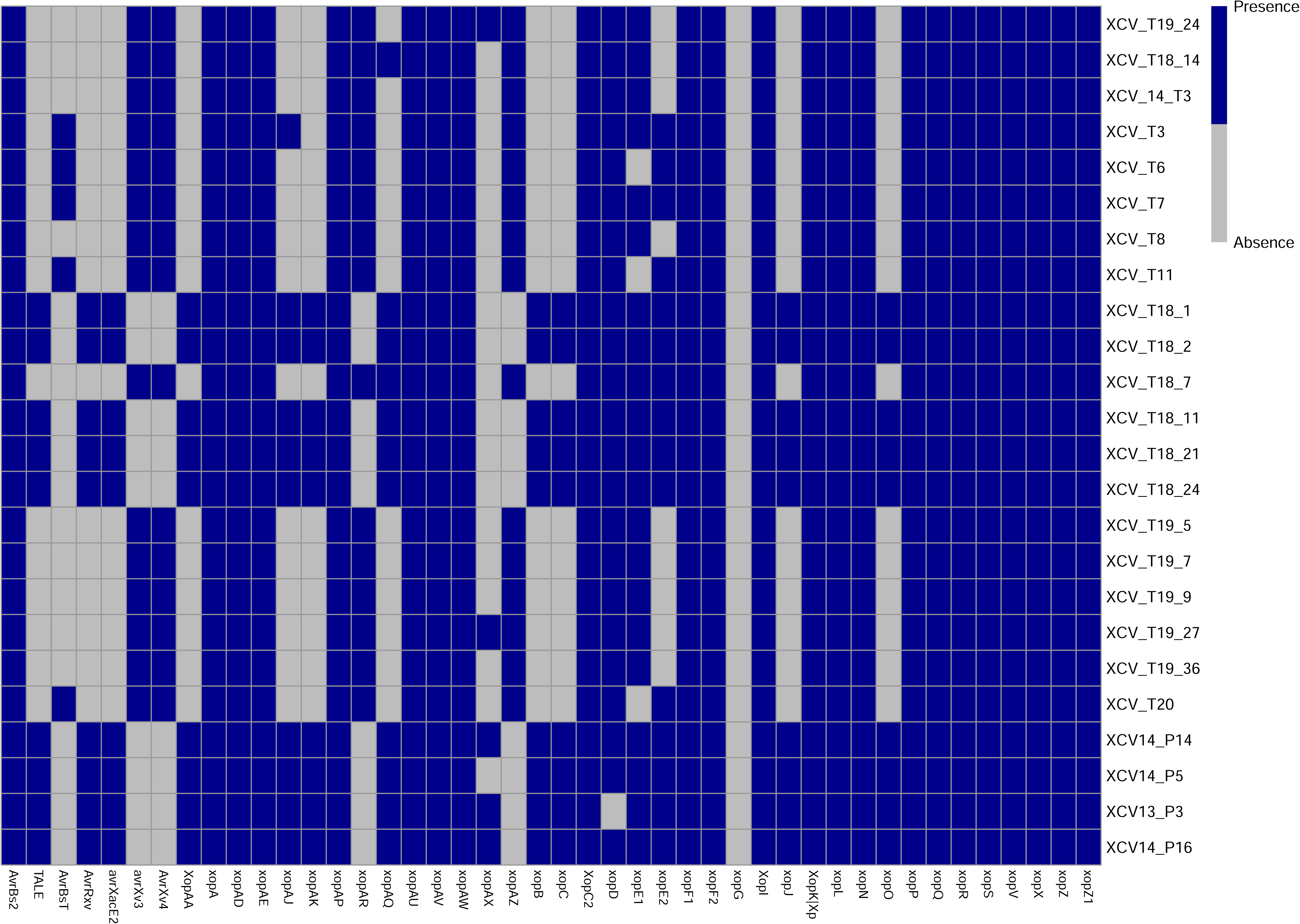
Effector profiles of sequenced *Xee* and *Xep* strains based on BLAST search. All the strains possess the bacterial spot xanthomonads core effectors.

## Discussion

Seasonal crop production requires growers to make timely decisions on plant disease management. The urgency and potential impact of this decision becomes more important during unexpected plant disease outbreaks. In this study we characterized the phenotypic and genomic profiles of strains isolated in Oklahoma following an outbreak of tomato bacterial spot in 2018 and 2019 as well as a few strains isolated in previous years. The whole genome comparisons and phylogeny based on core genes identified *Xee* and *Xep* as the two species causing bacterial spot of tomato and pepper in east and central Oklahoma. Whole genome comparisons also identified *X. arboricola, Xanthomonas alfalfae* subsp. *citrumelonis* and a potentially new species. These three strains, isolated from tomato, were not pathogenic and produced HRs on the tomato cultivar Bonny Best. Based on ANI and is-DDH values, strain XCV_T1 belongs to a new species that we have named *X. oklahomensis* sp. nov. The species description is provided below. Further investigation of non-bacterial spot xanthomonads in the pathogenesis and epidemiology of bacterial spot pathogens will be required to determine their role in the microbiome of bacterial spot disease (Sadhukhan et al., 2024).

We identified three tomato races, T1, T3 and T4, that cause bacterial spot of tomato in Oklahoma, with T4 strains shown to cause disease on pepper. Currently, strains of *Xanthomonas* species that cause bacterial spot disease on tomato are assigned into five races (Stall *et al*. 2009; Jibrin *et al*. 2022). Strains belonging to T1 race are typically *Xee* while T3, T4 and T5 races are *Xep* (Stall *et al*. 2009; Jibrin *et al*. 2022). In our study, strains isolated in 2018 consisted of T1 and T3 strains, while strains isolated in 2019 were exclusively T4 strains. The change in tomato races over just two seasons is striking. While race shifts within a single year have been reported frequently in pepper production systems, race shifts in tomato production systems tend to be more gradual, over a more sustained period (Klein-Gordon *et al*. 2021; Rotondo *et al*. 2022; Chen *et al*. 2024; Subedi *et al*. 2023). For example, the previous study on pepper bacterial strains in Oklahoma identified rapid shift in race population that overcame *Bs2* resistance in pepper in a single season (Damicone *et al*. 2021). Similarly, race shifts in pepper production system within a growing season have been documented previously in Florida and North Carolina (Pohronezny *et al*. 1992; Kousik and Ritchie, 1996). In Florida tomato production system, the first T4 strain was identified in 1998, seven years after detecting T3 strains (Wang *et al*. 1990; Jones *et al*. 1995; Astua-Monge *et al*. 2000). By 2012, about 15 years after the emergence of T4 strains, the T4 strains displaced T3 strains from tomato fields (Horvath *et al*. 2012). The identification of only strains of T4 race within a year of recovering T1 and T3 races during this outbreak in Oklahoma tomato production system is therefore striking. Interestingly, in this study, tomato farms in Atoka, Hughes, Payne and Rogers counties were sampled twice in 2018 and 2019, with some of the surveys repeated on the same farms in both years. In general, Oklahoma tomatoes are produced in small acreages, mostly for farmers’ markets, and the same field can be used for growing different kinds of other crops. A rapid replacement could be explained by lack of overwintering of the pathogen population, a change in source of planting material, or competition between strains of different genotypes.

At the scale of genotype competition, it took bacteriocin-producing *Xep* strains about 15 years to completely displace *Xee* strains in Florida tomato production system (Hert *et al*. 2005; Tudor-Nelson *et al*. 2003; Horvath *et al*. 2012; Klein-Gordon *et al*. 2021). Similarly, in Taiwan, almost a decade elapsed since the introduction of *Xep* before *Xee* was completely displaced from tomato production fields (Chen *et al*. 2024). In Brazil, survey of the bacterial spot complex on tomatoes over a four-year period identified a single *Xee* strain in the first year in one county, alongside *Xep* T3 strains (Araujo *et al*. 2016). Although subsequent surveys did not include the county and inferences on species displacement in that county cannot be confirmed, surveys in the next three years did not recover *Xee* strain from other counties (Araujo *et al*. 2016). In this study however, while *Xee* strains were recovered in 2018, no *Xee* strain was recovered in 2019 from the same survey area. Strains of *Xep* have been shown to produce bacteriocins against *Xee* which, over time, resulted in the now dominant *Xep* in Florida tomato production fields (Hert *et al*. 2005; Klein-Gordon *et al*. 2021). The displacement of *Xep* T3 by T4 strains has been explained in terms of fitness conferred by effector content variation, rather than the production of bacteriocins (Timilsina et al. 2016). Combined with previous observations of race shifts in a growing season in bacterial spot of pepper (Pohronezny *et al*. 1992; Kousik and Ritchie, 1996; Damicone *et al*. 2021), our results suggest that rapid changes in races and, potentially, different genotypes of the bacterial spot pathogen may be more common than previously thought, presenting a challenge for disease management. Alongside bacterial competition, transplants and weather events such as hurricanes can contribute to the spread of pathogens, including *Xanthomonas* species that can impact species composition (Bock *et al*. 2010; Abrahamian *et al*. 2021). Future studies to improve understanding will include elucidating the nature of bacteriocin production in these strains as well as other factors such as tornadoes, planting materials and pathogen life cycle that may be unique to Oklahoma tomato production system and contribute to rapid changes in population within and between production seasons.

All the strains in this study were sensitive to streptomycin. We are not aware of previous reports of the use of streptomycin in managing bacterial spot disease in Oklahoma, so sensitivity to streptomycin is not unexpected as the antibiotic may not be widely used. However, streptomycin resistance is present in *Xep* in Florida (Klein-Gordon *et al*. 2021). Most of the strains were sensitive to copper sulfate treatment, but a few strains were tolerant. If the copper tolerant strains contain plasmid-borne copper resistance genes, *copLAB*, there is the possibility of spread of copper resistance through horizontal gene transfer (Jibrin *et al*. 2024; Bibi *et al*. 2024; Kaur *et al*. 2024). In Florida, copper resistance genes borne on both conjugative plasmid and in the chromosome are responsible for widespread copper resistance in *Xep* (Bibi *et al*. 2024; Kaur *et al*. 2024). Copper resistance in a bacterial spot population can be variable and can render the chemical control of bacterial spot of tomato difficult (Khanal *et al*. 2020; Abbasi *et al*. 2015; Abrahamian *et al*. 2019; Adhikari *et al*. 2019; Jibrin *et al*. 2024; Bibi *et al*. 2024; Kaur *et al*. 2024). Since copper-based bactericides remain the major chemical control of choice for growers, monitoring the bacterial spot population for copper resistance and deployment of integrated management approaches that reduce the use of copper could be important in helping growers manage the disease.

The *Xee* strains in this study exhibited differences in amylolytic activity. While *Xee* strains from tomato were amylolytic, strains from pepper were non-amylolytic. Historically, *Xee* strains are primarily known to be non-amylolytic and this characteristic was used among the preliminary tools to differentiate them from the amylolytic *Xep* (Jones *et al*. 2004). However, recent analyses of *Xee* strains from Florida identified a genetic lineage associated with amylolytic activity and possess an intact alpha-amylase gene as found in *Xep* (Subedi *et al*. 2024b). Amylolytic *Xee* strains were also identified in strains from Ohio, Taiwan, Mexico and China (Jones *et al*. 2004; Subedi *et al*. 2024b). The specific advantage of extracellular amylase production remains unclear and requires further elucidation (Subedi *et al*. 2024b). However, monitoring this emergent phenotype in *Xee* will provide added insights into population diversity and change over time.

Type III secretion system (T3SS) effectors are key proteins in host-pathogen interactions and pathogenicity of bacterial spot xanthomonads on their hosts (White *et al*. 2009; Potnis *et al*. 2011; Schwartz *et al*. 2015). All the sequenced *Xee* and *Xep* strains harbor all previously described core T3SS effectors that are found in all bacterial spot xanthomonads (Potnis *et al*. 2011, Schwartz *et al*. 2015). We propose that the effector XopAU be added to the core effectors of bacterial spot xanthomonads since they are now frequently reported to be present in all recently published genomes of bacterial spot xanthomonads, especially *Xee*, *Xep* and *X. hortorum* pv. *gardneri* (Jibrin *et al*. 2024; Subedi *et al*. 2024a; Subedi *et al*. 2024b; Timilsina *et al*. 2025). Furthermore, our results showed that although all the pre-2018 tomato *Xep* strains except strain XCV_T7 possess the effector AvrBsT, none of the 2018 and 2019 strains possess this effector gene. Loss of the effector gene *avrBsT* was previously associated with host expansion of some strains to pepper but it was also shown that strains that lacked *avrBsT* do not necessarily cause disease on pepper (Schwartz *et al*. 2015; Newberry *et al*. 2018). Similarly, while some studies also suggested the role of the TAL effector, AvrHah1, in promoting the pathogenicity of *Xep* on pepper, tomato *Xep* strains without AvrHah1 can equally be pathogenic on pepper (Newberry *et al*. 2018; Subedi et al. 2024a). None of the *Xep* T4 strains with expanded host range to pepper has any hit for a TAL effector, suggesting that factors other than TALEs have been responsible for the expanded host range as previously suggested. Although genome-wide association studies attempted to narrow down candidate genes responsible for host expansion of *Xep* to pepper, several factors are still needed to be elucidated (Newberry *et al*. 2023).

## Conclusions and future recommendations

In conclusion, this study contributes to understanding seasonal population shifts in bacterial plant pathogens and further lays a foundation for monitoring and management of the bacterial spot of tomato and pepper pathogen in Oklahoma. We identified the apparent displacement of *Xee* by *Xep* and a shift from T3 to T4 race within two years in Oklahoma. These findings, alongside seasonal race flux in *Xee* on pepper reported earlier (Pohronezny *et al*. 1992; Kousik and Ritchie, 1996; Damicone *et al*. 2021), suggest that monitoring the population of the pathogens causing bacterial spot disease on tomato and pepper during the production season and at annual intervals may be critical to improved management decisions. Our studies further demonstrate the expansion of host range of *Xep* to pepper hosts, confirming recent reports (Newberry *et al*. 2018; Subedi *et al*. 2024a) Additional field surveys will be required to confirm the current population of bacterial spot strains on tomato and pepper in Oklahoma and associated genomic evolution. Future studies exploring the progressive evolution of historical strains and newly recovered strains will be required to provide adequate management recommendations to growers.

### Xanthomonas oklahomensis sp. nov

The characteristics are as described for the genus (Vauterin *et al*. 1995). The type strain is XCV_T1 (also XCVT1). Colonies are mucoid yellow on nutrient agar medium and can grow at 28 ^0^C. ANI between XCV_T1 and the *Xanthomonas* type strains placed XCV_T1 closest to *X. hydrangea* at 91% similarity. Based on dDDH, *X. oklahomensis* is closest to *X. cynarae* CFBP4188 and *X. hydrangea* with 67.6% and 66.2% similarity, respectively. *X. oklahomensis* is neither pathogenic on tomato nor pepper. The type strain is sensitive to streptomycin and copper at 200 ppm. Strain XCV_T1 lacks copper resistance genes but is amylolytic and has an intact alpha amylase gene. The NCBI taxonomy ID is 3391446. The NCBI accession number is JBLDYR000000000.

## Funding Acknowledgement

This research was made possible in part by funding through Oklahoma Agricultural Experimental Station start-up HATCH Grant to Dr. Mustafa O. Jibrin.

