## Supplementary File 1 for "Rapid population flux in bacterial spot xanthomonads during a transition in dominance between two genotypes in consecutive tomato production seasons and identification of a new species *Xanthomonas oklahomensis* sp. nov"

**S1. Survey and preliminary characterization of recovered strains**

**S1.1** Occurrence and severity of bacterial spot disease on tomato in Oklahoma

| County^a^ | Year | Disease Incidence^b^ | Defoliation^c^ |
| --- | --- | --- | --- |
| Payne (n=2) | 2018 | 31.7 | 6.7 |
| Oklahoma (n=1) | 2018 | 26.7 | 10 |
| Rogers (n=1) | 2018 | 3.3 | 0.0 |
| Canadian (n=3) | 2018 | 46.7 | 15.6 |
| Pottawatomie (n=1) | 2018 | 13.3 | 6.7 |
| Atoka (n=1) | 2018 | 10 | 0 |
| Hughes (n=1) | 2018 | 46.7 | 13.3 |
| Logan (n=1) | 2018 | - | - |
| Mean |  | 25.5 | 7.5 |
| Payne (n=2) | 2019 | 20.8 | 3.3 |
| Rogers (n=2) | 2019 | 56.7 | 29.2 |
| Hughes (n=1) | 2019 | 40 | 3.3 |
| Atoka (n=2) | 2019 | 81.7 | 36.7 |
| Bryan (n=2) | 2019 | 43.3 | 28.3 |
| Pontotoc (n=1) | 2019 | 63.3 | 6.7 |
| Mean |  | 51 | 17.9 |

^a^ County where bacterial spot disease was diagnosed. The survey also included Septoria leaf spot caused by *Septoria lycopersici* (in Rogers and Cherokee counties in 2018) and early blight caused by *Alternaria solani* (in Bryan County in 2019) but this is not shown in this table.	The “n” in parenthesis after each county shows the number of field sites confirmed for bacterial spot in each county. Disease diagnosed by culturing of yellow-mucoid colonies typical of *Xanthomonas* spp. from symptomatic leaves on nutrient agar and PCR detection of partial *hrcN* DNA fragment conserved among *Xanthomonas* by PCR-assay.

^b^ Disease Incidence defined as the percentage of leaves with symptoms of bacterial spot, was visually assessed on three plants / site.

^c^ Defoliation was visually assessed on three plants /site. Defoliation was identified as axils with leaf abscission (no attached petioles) and was expressed as a percentage of total number of leaves, including defoliated leaves.

**S1.2.** Identification of *Xanthomonas* isolates from tomatoes in Oklahoma collected between 1998 and 2014 based on partial sequence of *hrcN* gene

| Isolate | County^a^ | Year | NCBI Accession Number | Species identification^b^ |
| --- | --- | --- | --- | --- |
| XCVT1 (XCV_T1) | Tulsa | 1998 | PV430080 | *X. euroxanthea^c^* |
| XCVT2 | Tulsa | 2001 | PV430043 | *Xep^d^* |
| XCVT3 (XCV_T3) | Oklahoma | 2001 | PV430031 | *Xep* |
| XCVT4 | Oklahoma | 2002 | PV430044 | *Xep* |
| XCVT5 | Oklahoma | 2002 | PV430045 | *Xep* |
| XCVT6 (XCV_T6) | Oklahoma | 2002 | PV430046 | *Xep* |
| XCVT7 (XCV_T7) | Payne | 2002 | PV430047 | *Xep* |
| XCVT8 (XCV_T8) | Payne | 2002 | PV430048 | *Xep* |
| XCVT9 | Tulsa | 2002 | PV430032 | *Xep* |
| XCVT11 (XCV_T11) | Canadian | 2002 | PV430051 | *Xep* |
| XCVT14-T3 | Tulsa | 2002 | PV430079 | *X. vesicatoria* |
| XCVT15 | Tulsa | 2002 | PV430052 | *Xep* |
| XCVT17 | Tulsa | 2002 | PV430053 | *Xep* |
| XCVT18 | Oklahoma | 2004 | PV430054 | *Xep* |
| XCVT19 | Oklahoma | 2004 | PV430055 | *Xep* |
| XCVT20 (XCV_T20) | Oklahoma | 2004 | PV430056 | *Xep* |

^a^ County in Oklahoma where a *Xanthomonas* species was isolated from tomatoes.

^b^ Strains were identified by PCR amplification of partial region of the *hrcN* gene using RST65/RST69 primers followed by Sanger sequencing of purified PCR products (Obradovic *et al.* 2004).

^c^ Genome sequence identified a novel species, *Xanthomonas oklahomensis* sp. nov.

^d^*Xep=* *X. euvesicatoria* pv. *perforans*

**S1.3.** Identification of *Xanthomonas* strains from tomatoes in Oklahoma in 2018 and 2019^a^

| Isolate | County^b^ | NCBI Accession number | Species Identification^c^ |
| --- | --- | --- | --- |
| **2018** |  |  |  |
| XCVT18-1 (XCV_T18_1) | Payne | PV430025 | *Xee^c^* |
| XCVT18-2 (XCV_T18_2) | Payne | PV430026 | *Xee* |
| XCVT18-6 | Oklahoma | PV430033 | *Xep^d^* |
| XCVT18-7 (XCV_T18_7) | Rogers | PV430034 | *Xep* |
| XCVT18-8 | Rogers | PV430035 | *Xep* |
| XCVT18-9 | Rogers | PV430036 | *Xep* |
| XCVT18-11 (XCV_T18_11) | Payne | PV430027 | *Xee* |
| XCVT18-12 | Canadian | PV430028 | *Xee* |
| XCVT18-13 | Canadian | PV430037 | *Xee* |
| XCVT18-14 (XCV_T18_14) | Canadian | PV430049 | *Xee* |
| XCVT18-15 | Canadian | PV430038 | *Xep* |
| XCVT18-16 | Pottawatomie | PV430039 | *Xep* |
| XCVT18-18 | Atoka | PV430029 | *Xee* |
| XCVT18-21 (XCV_T18_21) | Logan | PV430081 | *X. arboricola* pv. *pruni* |
| XCVT18-24 (XCV_T18_24) | Logan | PV430030 | *Xee* |
| **2019** |  |  |  |
| XCVT19-1 (XCV_T19_1) | Payne | PV430082 | *X. arboricola* pv. *juglandis* |
| XCVT19-5 (XCV_T19_5) | Payne | PV430057 | *Xep* |
| XCVT19-6 | Payne | PV430058 | *Xep* |
| XCVT19-7 (XCV_T19_7) | Payne | PV430059 | *Xep* |
| XCVT19-8 | Payne | PV430060 | *Xep* |
| XCVT19-9 (XCV_T19_9) | Payne | PV430061 | *Xep* |
| XCVT19-11 (XCV_T19_11) | Payne | PV430062 | *Xep* |
| XCVT19-18 | Rogers | PV430063 | *Xep* |
| XCVT19-19 | Rogers | PV430064 | *Xep* |
| XCVT19-20 | Rogers | PV430065 | *Xep* |
| XCVT19-21 | Rogers | PV430066 | *Xep* |
| XCVT19-23 | Hughes | PV430040 | *Xep* |
| XCVT19-24 (XCV_T19_24) | Atoka | PV430067 | *Xep* |
| XCVT19-26 | Atoka | PV430068 | *Xep* |
| XCVT19-27 (XCV_T19_27) | Atoka | PV430069 | *Xep* |
| XCVT19-28 | Atoka | PV430070 | *Xep* |
| XCVT19-29 | Atoka | PV430071 | *Xep* |
| XCVT19-30 | Atoka | PV430072 | *Xep* |
| XCVT19-31 | Atoka | PV430050 | *Xep* |
| XCVT19-35 | Bryan | PV430041 | *Xep* |
| XCVT19-36 (XCV_T19_36) | Bryan | PV430073 | *Xep* |
| XCVT19-37 | Bryan | PV430074 | *Xep* |
| XCVT19-38 | Pontotoc | PV430075 | *Xep* |
| XCVT19-39 | Pontotoc | PV430076 | *Xep* |
| XCVT19-40 | Pontotoc | PV430077 | *Xep* |
| XCVT19-41 | Pontotoc | PV430042 | *Xep* |
| XCVT19-42 | Pontotoc | PV430078 | *Xep* |

^a^ Strains were identified by PCR amplification of partial region of the *hrcN* gene using RST65/RST69 primers followed by Sanger sequencing of purified PCR products (Obradovic *et al.* 2004).

^b^ County in Oklahoma where a *Xanthomonas* species was isolated from tomato.

^c^*Xee* is *X. euvesicatoria* pv. *euvesicatoria*

^d^*Xep* is *X. euvesicatoria* pv. *perforans*
