## Supplementary File 3 for "Rapid population flux in bacterial spot xanthomonads during a transition in dominance between two genotypes in consecutive tomato production seasons and identification of a new species *Xanthomonas oklahomensis* sp. nov"

### Slide 1
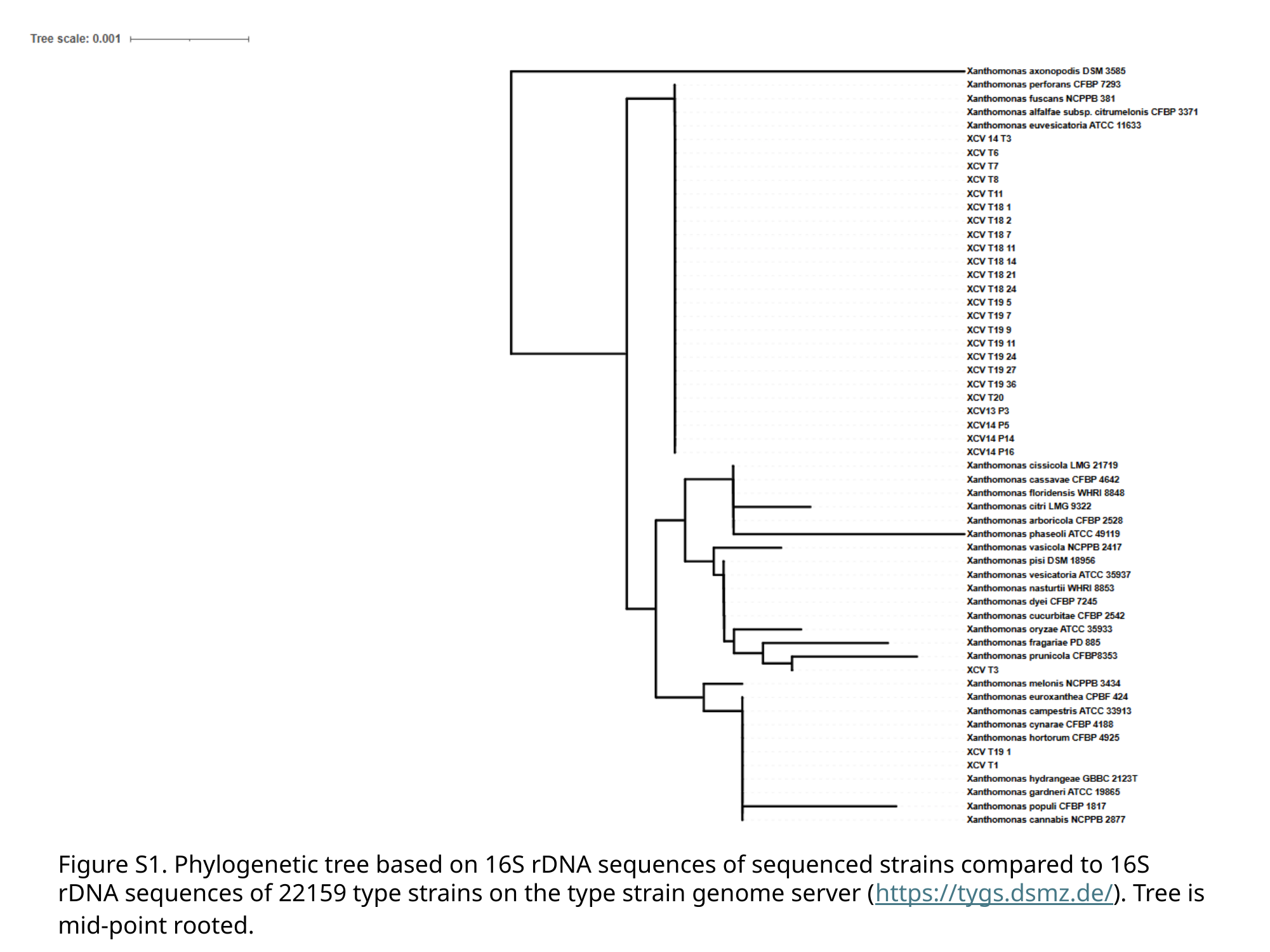

Figure S1. Phylogenetic tree based on 16S rDNA sequences of sequenced strains compared to 16S rDNA sequences of 22159 type strains on the type strain genome server (https://tygs.dsmz.de/). Tree is mid-point rooted.

### Slide 2
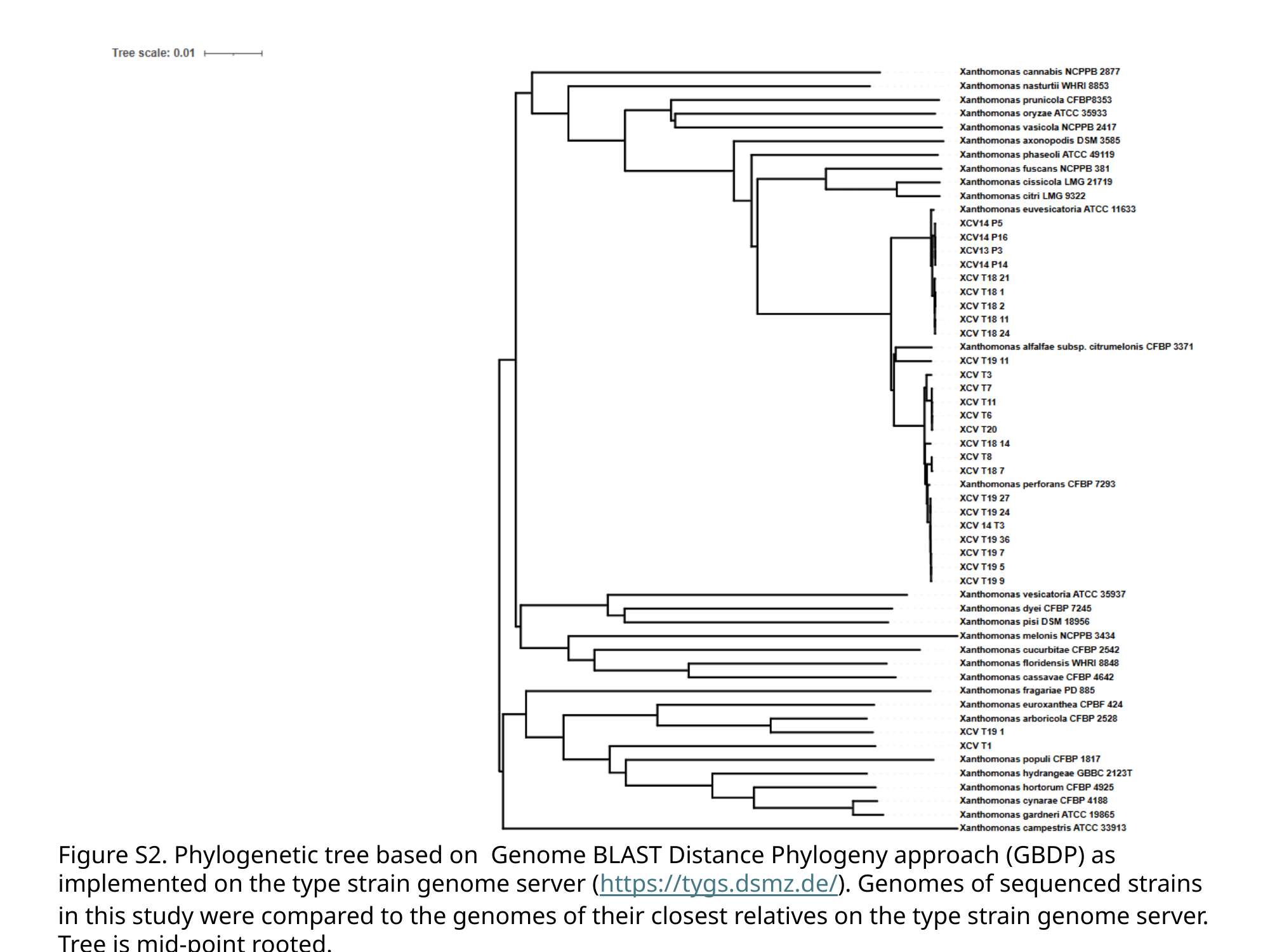

Figure S2. Phylogenetic tree based on Genome BLAST Distance Phylogeny approach (GBDP) as implemented on the type strain genome server (https://tygs.dsmz.de/). Genomes of sequenced strains in this study were compared to the genomes of their closest relatives on the type strain genome server. Tree is mid-point rooted.

### Slide 3
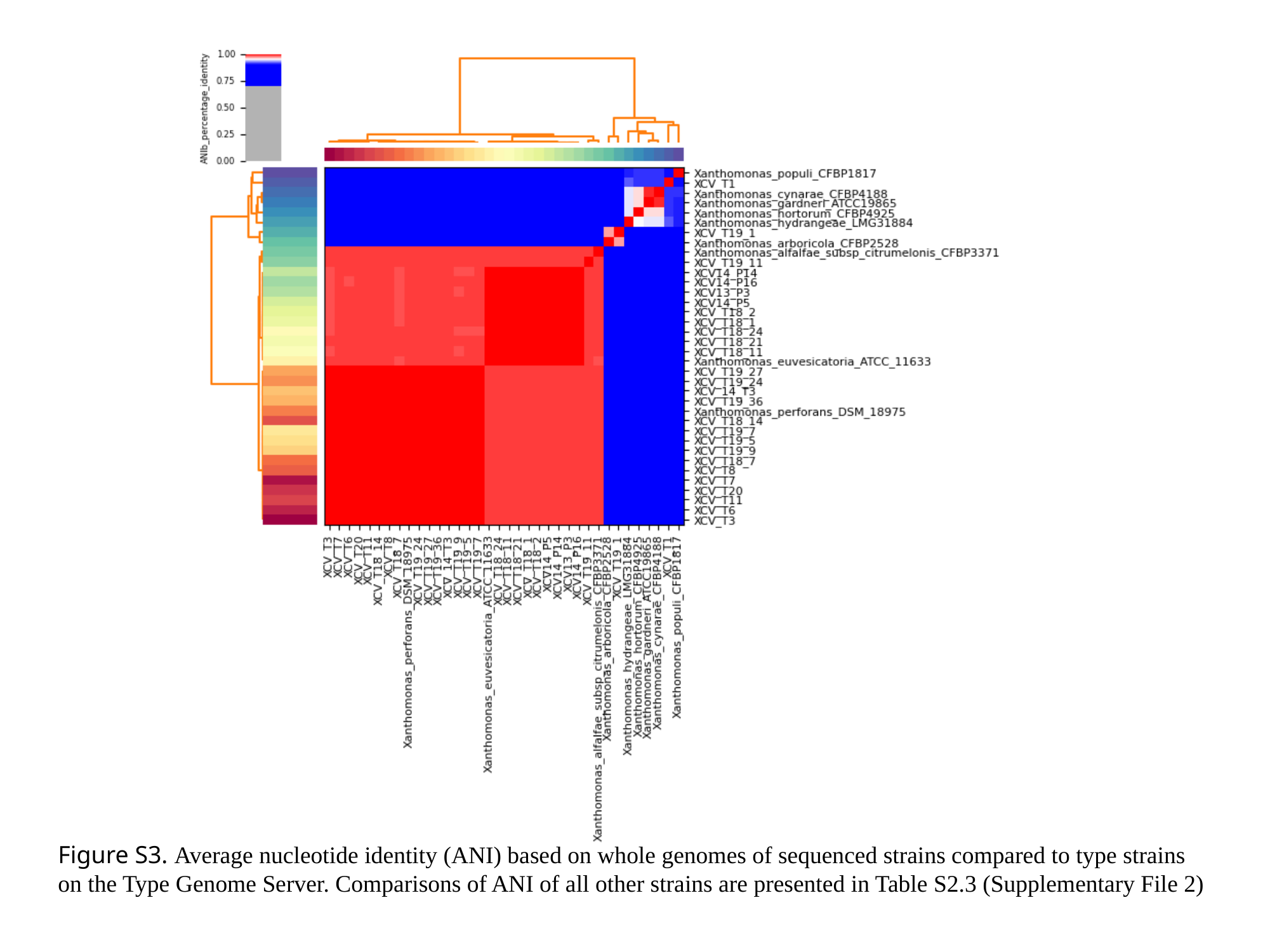

Figure S3. Average nucleotide identity (ANI) based on whole genomes of sequenced strains compared to type strains on the Type Genome Server. Comparisons of ANI of all other strains are presented in Table S2.3 (Supplementary File 2)
